# Real-time markerless video tracking of body parts in mice using deep neural networks

**DOI:** 10.1101/482349

**Authors:** Brandon Forys, Dongsheng Xiao, Pankaj Gupta, Jamie D Boyd, Timothy H Murphy

**Author notes:** **Correspondence should be addressed to**: Timothy H Murphy Address: 2255 Wesbrook Mall, Detwiller Pavilion, Vancouver, B.C. V6T 1Z3, Canada. Co-first authors (Brandon Forys, Dongsheng Xiao).

## Abstract

Markerless and accurate tracking of mouse movement is of interest to many biomedical, pharmaceutical, and behavioral science applications. The additional capability of tracking body parts in real-time with minimal latency opens up the possibility of manipulating motor feedback, allowing detailed explorations of the neural basis for behavioral control. Here we describe a system capable of tracking specific movements in mice at a frame rate of 30.3 Hz. To achieve these results, we adapt DeepLabCut – a robust movement-tracking deep neural network framework – for real-time tracking of body movements in mice. We estimate paw movements of mice in real time and demonstrate the concept of movement-triggered optogenetic stimulation by flashing a USB-CGPIO controlled LED that is triggered when real time analysis of movement exceeds a pre-set threshold. The mean time delay between movement initiation and LED flash was 93.44 ms, a latency sufficient for applying behaviorally-triggered feedback. This manuscript presents the rationale and details of the algorithms employed and shows implementation of the system using behaving mice. This system lays the groundwork for a behavior-triggered ‘closed loop’ brain-machine interface with optogenetic stimulation of specific brain regions for feedback.

## INTRODUCTION

Real-time movement tracking is a challenging computer vision problem that is crucial for constructing precise movement-triggered systems and brain-machine interfaces (BMIs). Most BMIs use electroencephalograms (EEGs) as inputs ^1^ to systems that provide feedback in near real time. We sought to explore methods by which we could precisely track movements of specific body parts in real time using computer vision techniques, to investigate whether video-based movement tracking could be used in BMIs.

Significant progress has been made in movement tracking and it is now possible to estimate and analyze poses in humans ^2^. Marker less, accurate tracking of specified movements without having to manually label large datasets as inputs for training ^3^ is possible. In particular, the approach presented by Mathis et al.’s (2018) “DeepLabCut” generalizes well across animals, and allows movement schemas to remain accurate across different mice. The flexibility offered by this approach is important for real-time movement tracking; instead of tracking movement based on databases of stereotyped movement data – such as those used by Insafutdinov et al. for pose estimation (2016). The DeepLabCut approach ^3^ generates models that can be more sensitive to the movements of animals under our specific laboratory conditions. Such sensitivity will help us overcome the limitations of a computer vision approach for measuring movement activity compared to more traditional BMI techniques, such as those utilizing EEG data ^1^ or single-neuron input data ^4^. Additionally, a robust, customizable framework for real-time tracking of specific body parts in atypical subjects (such as specific animals) would have many applications in psychiatry, rehabilitation engineering, and other fields that make use of BMIs. Our main motivation for developing a real-time movement tracking framework is to enable optogenetic ^5,6^ or sensory stimulation based on classification of movements of specific body parts. This framework may provide insight into the operations of the cortical regions involved in the coordination and planning of movements when used in combination with optogenetics^7–9^.

Although our aim is to employ optogenetics, we must first show that we can track the movement of specific body parts in real time. However, virtually all current real-time tracking systems used for animal tracking are based on blob detection algorithms ^10,11^ which, while more lightweight than pose estimation algorithms, are better suited to whole-body tracking than body part tracking. As such, while previous approaches to real-time tracking are effective for examining social interactions or holistic body movements, they are typically unable to discern small-scale movements – such as whisker or nose movements – which are crucial as inputs to an appropriately robust BMI. Therefore, here we discuss adaptations we have made to *DeepLabCut* ^3^, a precise movement tracking framework, in order to leverage it for real-time movement tracking and analysis on individual body parts in mice.

## MATERIALS AND METHODS

### Animals and surgery

Animal protocols (A13-0336 and A14-0266) were approved by the University of British Columbia Animal Care Committee and conformed to the Canadian Council on Animal Care and Use guidelines and animals were housed in a vivarium on a 12 h day light cycle (7 AM lights on). For head fixation experiments animals were anesthetized with isoflurane (2% in pure O_2_) and body temperature was maintained at 37°C using a feedback-regulated heating pad monitored by a rectal thermometer while they received a cranial window. Mice received an intramuscular injection of 40 μl of dexamethasone (2 mg/ml) and a 0.5 ml subcutaneous injection of a saline solution containing buprenorphine (2 μg/ml), atropine (3 μg/ml), and glucose (20 mm), and were placed in a stereotaxic frame. After locally anesthetizing the scalp with lidocaine (0.1 ml, 0.2%), the skin covering the skull was removed and replaced by dental cement ^7,12,13^. A metal screw was attached to the chamber for future head fixation during recordings. At the end of the procedure, the animal received a second subcutaneous injection of saline (0.5 ml) with 20 mM of glucose and recovered in a warmed cage for 30 min.

For movement recordings, the heads of awake mice were stabilized by attaching a skull-mounted screw to a pole mounted on a base-plate while the body was resting on a running wheel. We used a USB 3.0 webcam (Logitech BRIO, Logitech, Lausanne, Switzerland, 60 Hz) or a Raspberry Picam’s RGB sensor (Raspberry Pi Foundation, Cambridge, UK, 60 Hz) to capture body movements.

### Training movement tracking models

We use DeepLabCut ^3^ as the basis for our movement tracking framework. Using the standard protocol developed by Mathis et al. (https://github.com/AlexEMG/DeepLabCut), we train models to analyze movement of each mouse’s nose, left and right barrel, and left and right paws. All image processing, tracking, and LED output was carried out on a computer with 64 GB of RAM, 3.3 GHz and an Nvidia Titan Xp GPU; however, we also conducted a number of trials on another computer with 128 GB of RAM.

### Implementing real-time tracking

We examine two approaches to streaming video for real time tracking. In both approaches, we primarily investigate and collect data for the left and right paw movement; this is because, for the purposes of developing a robust real-time movement tracking paradigm, larger movements are easier to track. Additionally, the relatively high speed of paw movement enabled us to verify the fidelity of tracking. In the first approach, we stream a frontal video of the mouse (in the same position and under the same conditions as the videos on which the models were trained) *via* TCP, using a Raspberry Picam RGB sensor. We use a Python server script on the Raspberry Pi and a Python client script on the computer with DeepLabCut to accomplish this. In the second approach, we stream a video through the same neural network and video configuration as described above using a USB 3.0 webcam connected directly to the computer on which the DeepLabCut neural network model operates. Image retrieval, pre-processing, analysing and saving in serial loop is slow. To speed-up the real-time tracking, we split these serial operations into parallel tasks. An independent thread continuously captures the current frame from the camera and makes it available for pre-processing. Pre-processing and analysis processes run in serial and the results, along with the frame as passed to another independent thread that saves them. De-coupling 1) reading the frame from camera and 2) saving the frames and results from the main analysis process resulted in six-fold boost in the speed.

For each approach, we integrate our client script with DeepLabCut’s movement analysis functions. As each video frame arrives on the computer, we convert it to an 8-bit unsigned byte format and pass it to DeepLabCut’s pose analysis function. This function returns a set of six predicted locations on the body part being analyzed. Optionally, we then render these coordinates onto the newly analyzed frame using matplotlib (https://matplotlib.org/) and display the frame or save the frame. In the second approach, we shift the plot rendering process to a discrete thread in order to improve performance, and plot the coordinates using opencv2 (https://opencv.org/). We measure analysis performance through visual inspection and comparison of frame rates. The stream’s frame rate indicates the computational weight of the analysis; we optimize the computational environment in order to maximize the frame rate. We also manipulate lighting intensity and direction, camera position, stage position, resolution, and cropping: all of these factors have been observed to affect tracking accuracy.

### Prototyping optogenetic stimulation

In each approach, we calculate either a dynamic (cumulative standard deviation) or static threshold for movement classification. We then check if the average movement of the six points tracked on the selected body part exceeds this threshold.

In the first approach, if movement exceeds this threshold, we pass a packet containing “1” over UDP (over ethernet) to the Raspberry Pi, triggering the LED connected to the Raspberry Pi. If movement does not exceed this threshold, we extinguish the LED by sending a packet containing “0” over UDP (over ethernet) to the Raspberry Pi. In this approach, all LED operations are carried out asynchronously.

In the second USB 3.0 approach, if movement exceeds the threshold, we directly pass a command to turn the LED on or off over USB *via* an Adafruit FT232h breakout board (Adafruit Industries, New York, NY). In this approach, all LED operations are carried out asynchronously using a discrete thread.

#### Code Availability

We have released our modifications to DeepLabCut and pyftdi, which enable real-time pose estimation and LED testing, at https://github.com/bf777/DeepCutRealTime.

## RESULTS

We have evaluated several approaches for real-time tracking using DeepLabCut. The performance of each model was evaluated on frame rate over 120 seconds, through visual inspection, and through average likelihood of each prediction being correct. For the TCP approach using Raspberry Pi, the mean frame rate across all trials (*N* = 17) with frame recording was 1.73 Hz*, SD* = 0.10 Hz; the best performing model with frame recording was 2.00 Hz. For the wired approach using the USB 3.0 webcam, the mean frame rate across all trials (*N* = 37, 3 mice) with frame recording was 30.3 Hz, *SD* = 0.53 Hz. When LED feedback was not implemented, the mean frame rate across all trials (*N* = 3) without frame recording was 50.87 Hz, *SD* = 8.79 Hz; the best performing model under these conditions was 56.67 Hz.

We evaluated the performance of the optogenetic stimulation prototype by inspecting the delay between movement initiation and LED illumination in the behavioral video frames we recorded. Across all trials, the mean delay between movement initiation and LED illumination across trials (*N* = 37, 3 mice) was 93.44 ms, *SD* = 22.99 ms (Fig. 3, Supplementary video 1).

**Fig. 1.**
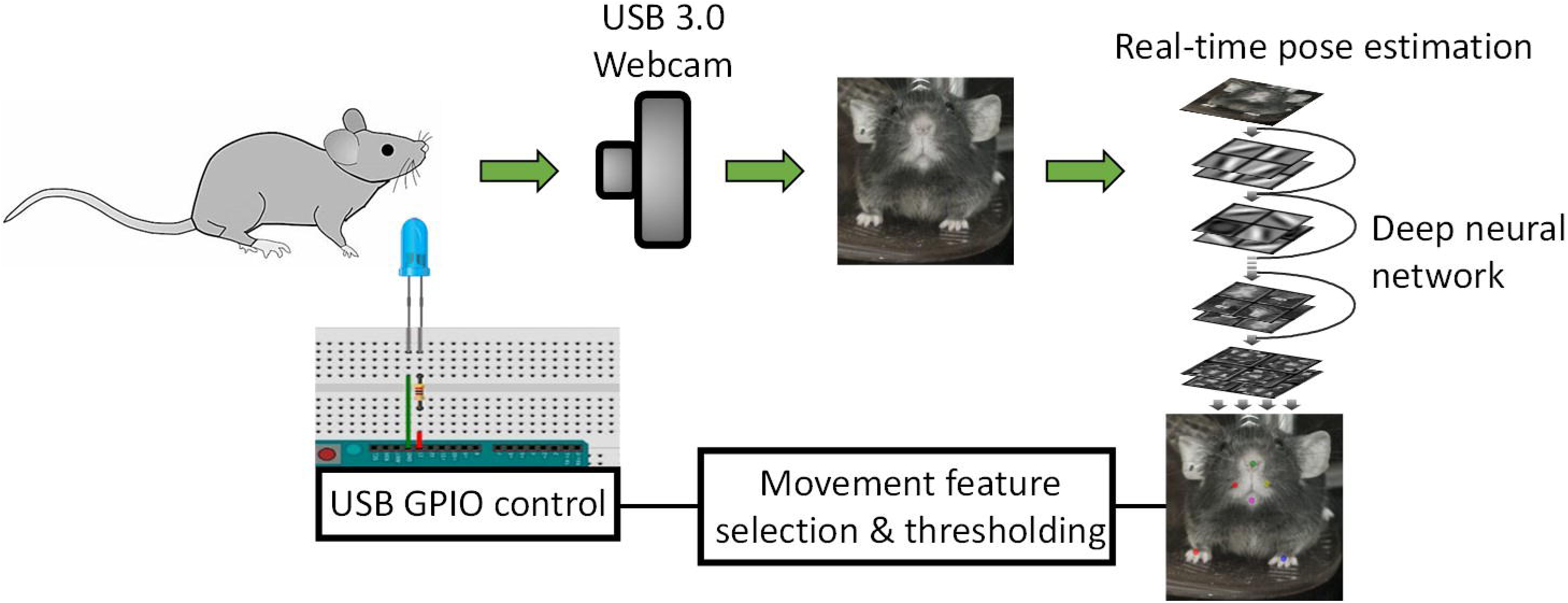
An overview of the video collection, pose estimation, and movement classification and feedback processes in our study.

**Fig. 2.**
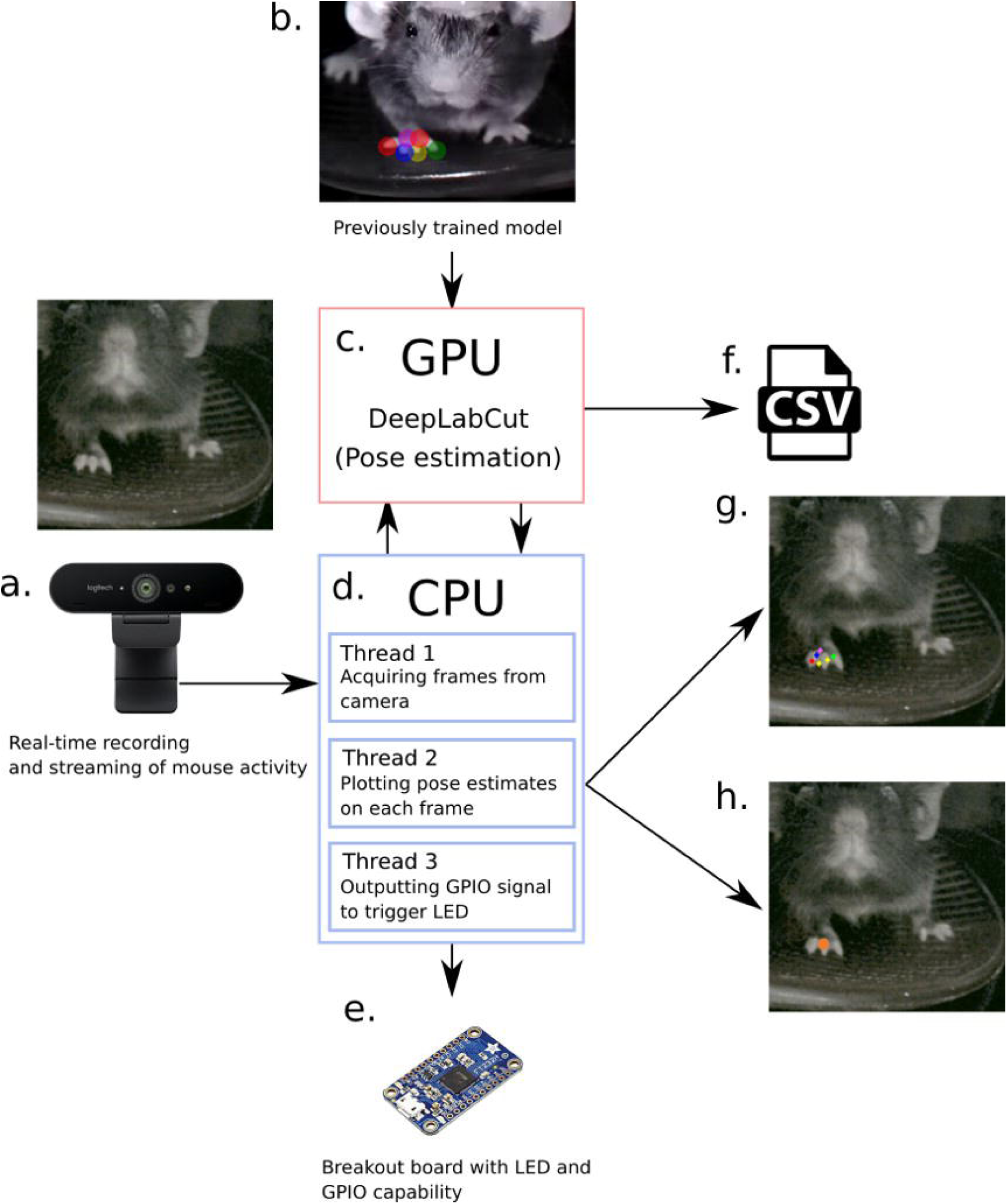
Overview of the parallel frameworks used to compute pose estimates and output a GPIO signal. a.) Mouse activity is recorded by webcam for 15 seconds. b.) A previously trained model of movement in the desired body part is selected. c.) Each frame of video from the webcam is sent over USB 3.0 to the computer; based on the previously trained video, *DeepLabCut* (operating on the GPU) estimates body part locations in each frame. d.) The estimated body part locations are then output to three CPU threads. The first thread collects each frame from the webcam and prepares it for processing. The second thread plots estimated body part locations on each frame before saving each frame – this step carries a negligible performance impact. If horizontal movement in a specified body part exceeds a pre-set threshold (defined as x: ≥20 px), the third thread is used to output a high signal output to a specified port on the breakout board using GPIO (e.). The LED connected to this port will then light up when the movement exceeds the threshold and be recorded by the behavioral video. Separately, all pose estimation data is saved to a CSV file (f.), and output frames can be set to render all points of tracking (g.) or an average of all points of tracking (h.).

**Fig. 3.**
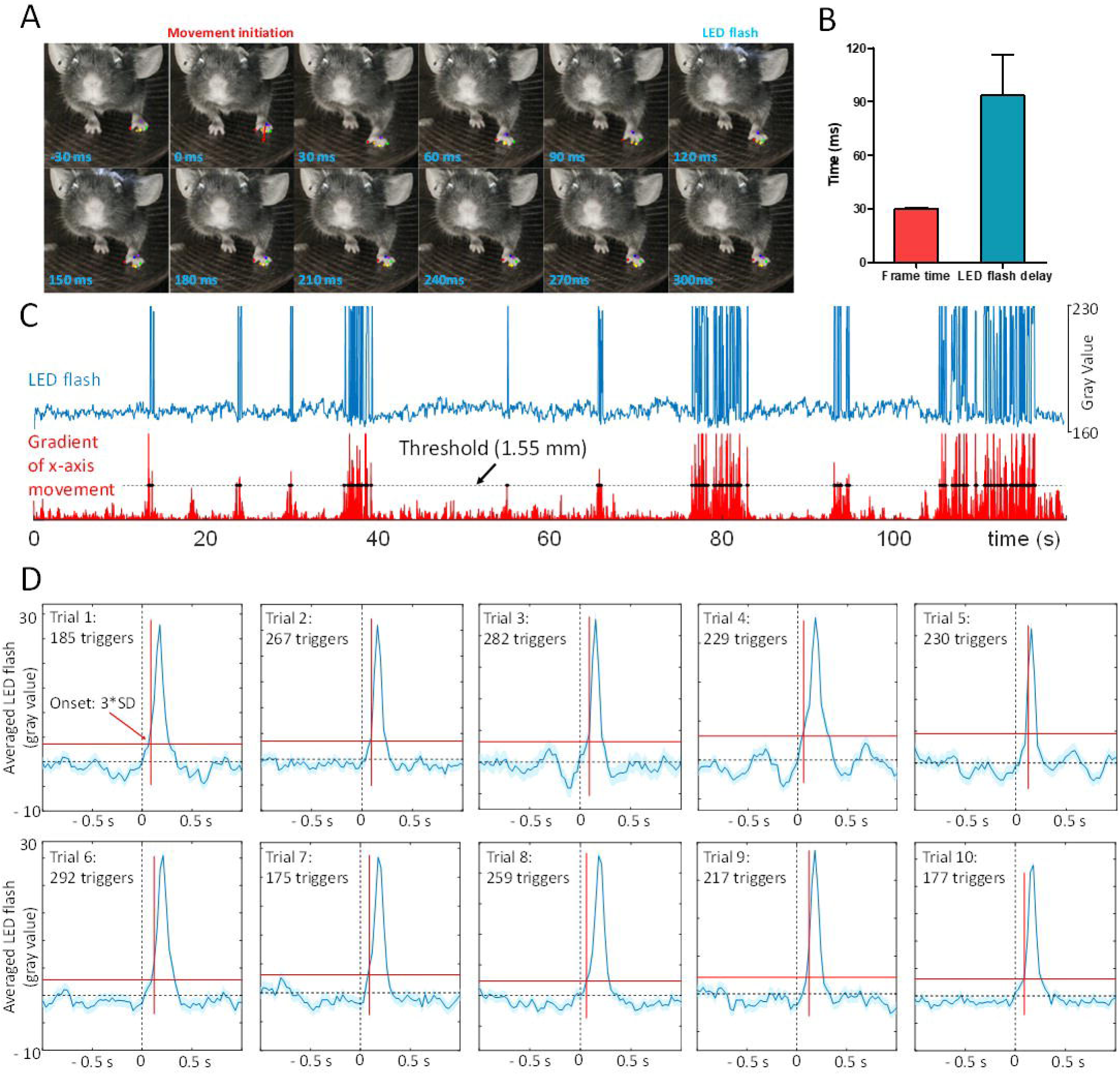
A). A visualization of the typical latency between movement initiation and the LED flash. B). The mean frame rate across all trials (*N* = 37, 3 mice) with frame recording was 30.3 Hz, *SD* = 0.53 Hz. the mean delay between movement initiation and LED illumination across trials (*N* = 37, 3 mice) was 93.44 ms, *SD* = 22.99 ms. C). LED flashes – as detected by changes in a region of interest on the region of the head where the LED flashes – are plotted against the x-axis movement on the left paw for an example run. The threshold is set as a difference of 12 px between the current frame and the last. D). GPIO output triggered average on LED flash of different experiments. The onset of LED flash is defined by 3*SD of baseline.

## DISCUSSION

We have adapted DeepLabCut for real-time tracking of behavior. We found that the USB 3.0 approach enabled a much higher frame rate than the TCP/UDP approach. Additionally, we were able to use threading to virtually eliminate any latency introduced by labelling frames. Shifting from matplotlib (Hunter, 2007) to native opencv2 (Bradski, 2000) plotting functions also appeared to contribute to the performance improvement. A number of factors could potentially affect the quality of the tracking. While tracking was typically robust, the most prominent of these factors was lighting; deviations from the lighting conditions of the videos on which we trained our models resulted in tracking of spurious body parts (such as the ear) or arbitrary points. In particular, regions of the video with high contrast relative to the intended body parts tended to be the focus for tracking when lighting conditions were incorrect. This may be a function of how the scoremap calculations that are involved in pose estimation are carried out within DeepLabCut ^3^. Additionally, fast running movements resulted in blurred body part tracking, which decreased quality; this is likely a function of the frame rate, which was influenced by 1) the GPU’s processing power, 2) the RAM of the computer, 3) the throughput of the connection between the camera and the computer (e.g. USB 3.0 vs. USB 2.0), and 4) the threading strategy used for image processing and LED control. As such, frame rate was improved by 1) upgrading the computer’s GPU and 2) upgrading its RAM, 3) switching from a USB 2.0 to a USB 3.0 webcam, and 4) switching from merely asynchronous plotting and LED operations to fully threaded versions of these operations. A smaller factor was the contrast level set on the webcam (likely for similar reasons to the effects of different lighting).

The relatively short delay between movement initiation and LED illumination is promising for further development of a movement-triggered biofeedback system with optogenetics. We have determined that, with only trivial modification, our existing mechanism can also interact with a trigger connected to a laser for optogenetic stimulation. However, this delay could be shortened by examining the hardware latency of our breakout board. The pyftdi library is one of the lowest-level interfaces between a computer and an LED; however, the breakout board is connected via USB 2.0, which presents a hardware bottleneck compared to the USB 3.0 technology used for our camera. As such, for future optogenetic work we need to account for and minimize this delay, perhaps by quantifying the delay and appropriately altering the frame rate to ensure that each LED flash is synchronized with a frame.

Further progress can likely be made by streamlining the deeper-level analytical operations of DeepLabCut and by further tweaking the threading strategy used. This would require deeper investigation into the most computationally intensive aspects of DeepLabCut; an especially important area to focus on would be parallelization of pose estimation operations in DeepLabCut. Additionally, it would be beneficial to batch the frames that are streamed into the pose estimation framework, in order to enable parallel processing of frames (which would then be sorted chronologically).

Our aim of integrating movement tracking with optogenetics will leverage recent advances in understanding of the mouse brain’s motor planning circuits^14^. Further directions for this research may combine movement tracking with two-photon microscopy to investigate whether real time DeepLabCut can be used to condition motor behaviours in mice through closed-loop feedback with the potential goal of understanding and localizing motor memory^15,16^.

### Conclusion

Our framework for real-time tracking based on DeepLabCut is capable of relatively high-speed tracking while maintaining performance. We demonstrate a movement magnitude classifier that can be used to trigger an LED, therefore prototyping a biofeedback brain-machine interface (BMI) that integrates optogenetic stimulation with movement tracking. This project forms the basis for future work on building a robust brain-machine interface that, through optogenetic stimulation that could be employed in forms of closed loop brain stimulation ^17,18^ and used to explore the function of various movement-related and somatosensory activities in the brain ^19^, including behavioural conditioning in mice via optogenetic stimulation ^20^. Such exploratory research could contribute to more advanced and effective BMIs that leverage both neural and non-neural data, adding greater diversity to the types of information that are integrated to treat somatosensory dysfunction.

**Video 1.** Real-time tracking and feedback LED flash. The time indicates the delay between movement initiation and LED flash in ms.

## Supporting information

## Acknowledgements

We thank Pumin Wang, Cindy Jiang for surgical assistance and technical assistance. We thank Hongkui Zeng and Allen Brain Institute for providing transgenic mice. We thank Rene Tandun for assisting in pose estimation model training.

## Contributions

B.F. and D.X. performed animal experiments. B.F., D.X., P.G., and J.D.B. developed and implemented the hardware and software for the apparatus. B.F. and D.X. wrote the analysis. B.F., D.X., and T.H.M. wrote the manuscript. B.F. and D.X. drew models and figures. D.X. and T.H.M. conceptualized the experiment.

